# AetherCell: A generative engine for virtual cell perturbation and *in vivo* drug discovery

**DOI:** 10.64898/2026.03.13.710968

**Authors:** Wenyuan Li, Yang Chen, Zhaoyi Peng, Lei Xiang, Dong Wang, Zhi Xie

**Affiliations:** State Key Laboratory of Ophthalmology, Zhongshan Ophthalmic Center, Sun Yat-sen University, Guangdong Provincial Key Laboratory of Ophthalmology and Visual Science, Guangzhou 410060, China; School of Basic Medical Sciences, State Key Laboratory of Southwestern Chinese Medicine Resources, Chengdu University of Traditional Chinese Medicine, Chengdu 611137, China

**Author notes:** Correspondence (Z.X.). These authors contributed equally (Wenyuan Li, Yang Chen).

## Abstract

Virtual cell modeling is currently hindered by a “data-utility paradox”: biological information is fragmented between context-rich clinical RNA-seq and perturbation-dense experimental assays, leading to poor predictive generalization in human contexts. Here, we introduce AetherCell, a generative foundation model that unifies these disparate domains into a shared, platform-aligned transcriptomic manifold. By implementing a specificity-driven learning framework, AetherCell successfully recovers low-frequency, mechanism-specific signals often obscured by systematic noise. Across extensive benchmarks, AetherCell demonstrates robust generalization, accurately predicting responses to unseen compounds and genetic perturbations. We show that the model effectively translates signals from simple cell lines to complex 3D organoids, achieving high-fidelity, whole-transcriptome prediction. Building on this foundational manifold, we demonstrate precise drug response prediction across biological scales—including patient-derived organoids and clinical cohorts. We also implement a phenotype-knowledge mixture-of-experts strategy for precision drug repurposing. This approach is validated by the *in vivo* discovery of teriflunomide for dry eye disease and dabigatran for ulcerative colitis. Together, AetherCell establishes a scalable, human-centric virtual cell framework for translational biology and accelerated drug discovery.

## Introduction

Non-animal, human-relevant evidence is increasingly shaping how new therapeutics are prioritized and de-risked^1^, intensifying interest in computational “virtual” systems that can predict how human tissues respond to chemical and genetic perturbations^2^. The effectiveness of such systems depends on their ability to model molecular interventions within human physiological environments. However, creating such models is currently hindered by a fundamental “data-utility paradox”: on the one hand, large public repositories contain hundreds of thousands of standardized human bulk RNA-seq profiles spanning diverse tissues and disease states, yet systematic perturbation annotations in these clinically grounded contexts remain scarce^3,4^. On the other hand, high-throughput screening platforms provide millions of perturbation signatures, but they are largely confined to a limited set of immortalized cancer cell lines, restricting direct translation into patient-relevant physiology^5,6^. A virtual cell model bridging these two regimes—dense perturbational coverage and rich human contextual diversity—enables scalable, human-centric virtual experimentation and is increasingly supported by regulatory openness to Non-Animal Methods (NAMs), such as the U.S. FDA Modernization Act 3.0^7^.

Although single-cell atlases have transformed cell-type-resolved biology, bulk RNA-seq datasets currently offer far deeper perturbational coverage at scale, capture robust population-level variation across thousands of clinical phenotypes, and serve as a practical standard for biomarker development and clinical transcriptomic readouts^3,4,10^. A principled bulk-first foundation could therefore act as a “universal coordinate system” in which perturbational effects learned from screens can be transferred into primary-like environments, organoid systems, and patient cohorts^11^.

Prior computational efforts have taken important steps toward this goal, from early pattern-matching frameworks to modern conditional and counterfactual generative models^5,6,12–14^. Yet three challenges repeatedly limit fidelity. First, platform and protocol discrepancies often dominate biological signals when transferring between landmark-gene assays and whole-transcriptome RNA-seq, leading to brittle representations^6,12^. Second, and more fundamentally, many models optimize toward deceptively strong global correlation by reproducing high-frequency “generic” responses (e.g., stress, metabolic shifts) that recur across diverse perturbations^15,16^. This “mean-state convergence” (a Type II failure) produces predictions that blur the mechanistic identity of a specific intervention, undermining the recovery of specific transitions in unseen, clinically grounded contexts^15^. Third, it remains unclear whether virtual cell models can faithfully predict behavior in complex human physiological contexts—such as patient-derived organoids or clinical cohorts—where the interplay of tissue architecture and individual variation often deviates significantly from the responses observed in standard *in vitro* models^17^.

Here we introduce AetherCell, a deep generative foundation model that unifies transcriptomic measurements into a shared, platform-aligned representation and enables transfer of perturbation effects into clinically grounded contexts. AetherCell learns a common coordinate system from large-scale RNA-seq and aligns high-throughput perturbation signatures to the same space, while conditioning generation on multi-modal priors capturing chemical structure and genetic regulatory logic. Crucially, we introduce a specificity-driven learning framework that suppresses non-specific stress-like responses, forcing the model to recover low-frequency mechanistic signals rather than generic averages.

Across comprehensive benchmarks, AetherCell preserves biological identity while bridging platform barriers, generalizing seamlessly from screened cell lines to independent RNA-seq perturbation datasets and complex 3D organoid environments. Building on this foundational manifold, we demonstrate two key translational applications. First, AetherCell-RP (AC-RP) enables high-fidelity prediction of drug response phenotypes, including therapeutic sensitivity in patient-derived organoids and clinical cohorts, drug combination effects, and genotype-linked vulnerabilities. Second, AetherCell-DR (AC-DR) facilitates precision drug repurposing by adaptively integrating transcriptomic reversal signals with structured knowledge priors. Finally, *in vivo* validation in two distinct disease models illustrates a practical, scalable pathway from virtual perturbation to actionable therapeutic hypotheses for human medicine.

## Results

### A unified generative foundation for cross-domain, multi-scale virtual cell modeling

Virtual cell modeling critically relies on the extensive perturbation datasets in human physiological contexts. To connect the rich human context of bulk RNA-seq data and the extensive perturbation profiles from L1000, we developed AetherCell, a deep generative framework that learns a unified transcriptomic manifold to enable the zero-shot transfer of perturbation effects from cell lines to complex human physiological environments (Fig. 1).

**Fig. 1.**
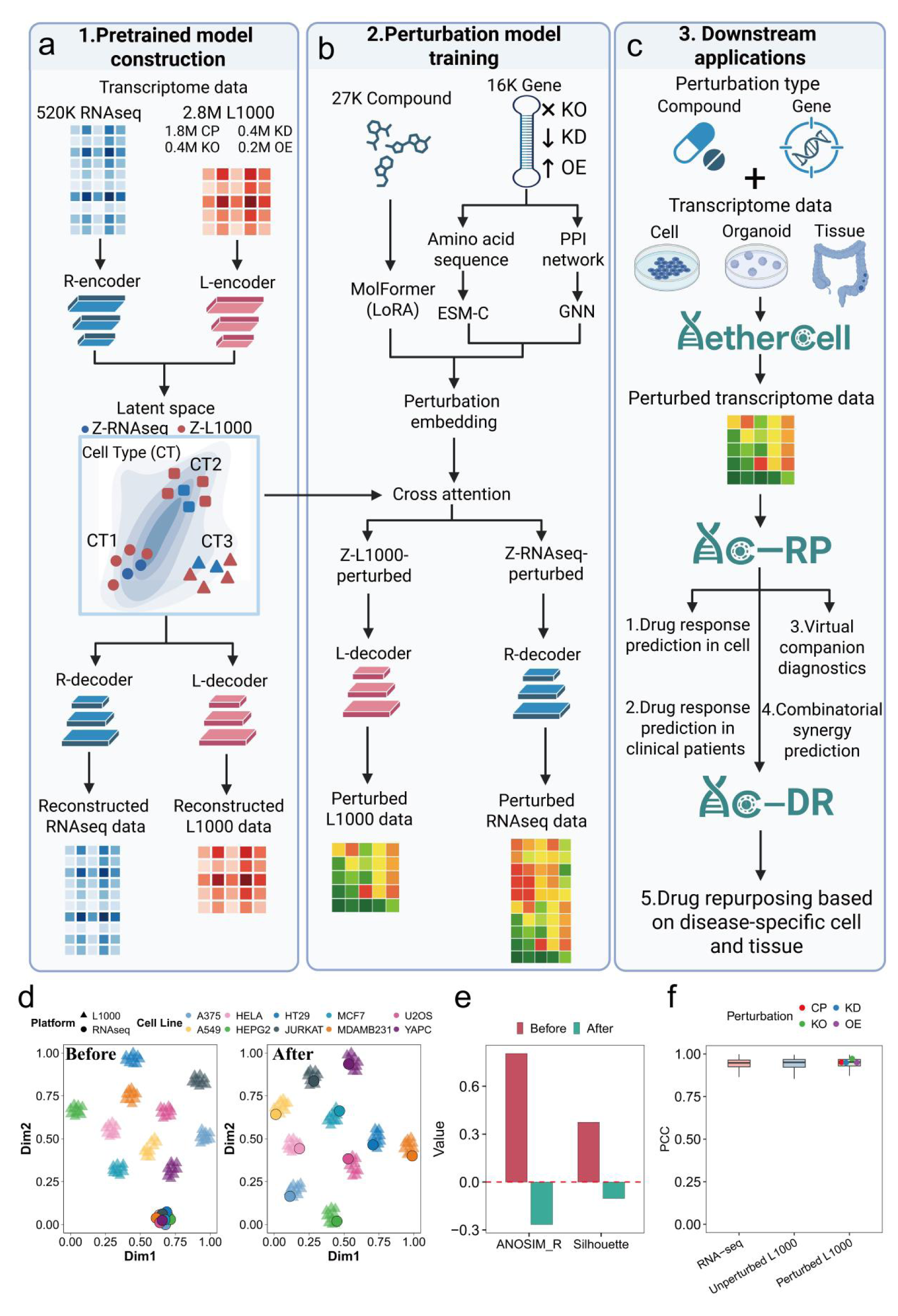
A unified generative framework for cross-platform perturbation modeling and zero-shot generalization. a, Construction of the unified transcriptomic manifold. A VAE learns a generalized latent space from human RNA-seq profiles (*n* = 519,609). Crucially, L1000 data (*n* = 2,811,324) are anchored to this manifold by aligning matched control samples to mitigate platform-specific batch effects. **b,** Integration of multi-modal foundation models. Small molecules (via MolFormer with LoRA) and gene perturbations (via ESM-C embeddings fused with STRING topology) are encoded to predict perturbation-specific latent shifts (Δ*z*) via cross-attention. **c,** Downstream applications. The unified representation supports tasks including drug sensitivity (IC50) and synergy prediction, companion diagnostics, and phenotype-based drug repurposing across cell lines, organoids, and tissues. **d, e,** Evaluation of platform alignment. (d) UMAP projections before and after alignment (Circles: RNA-seq; Triangles: L1000). (e) Alignment quantification via ANOSIM R and Silhouette scores on matched pairs (*n* = 125). Bars indicate metric values before (red) and after (green) processing. **f,** Reconstruction fidelity. Box plots showing the Pearson Correlation Coefficient (PCC) between reconstructed and ground-truth profiles for RNA-seq (*n* = 81,861); unperturbed L1000 (*n* = 28,304); and perturbed L1000 samples including Compound (CP, *n* = 270,885), Knockdown (KD, *n* = 67,976), Knockout (KO, *n* = 62,222), and Overexpression (OE, *n* = 19,750). Box plots indicate median, quartiles, and 1.5× IQR.

Our core strategy is first to construct a generalized transcriptomic manifold using a deep Backbone Variational Autoencoder (VAE)^18^ trained on the comprehensive RNA-seq corpus (Fig. 1a). Crucially, rather than treating L1000 data as a separate domain, we implemented a hierarchical Satellite-Backbone architecture to anchor the L1000 interface into this pre-established global manifold. This is achieved by explicitly aligning the latent representations of matched L1000 control samples to their RNA-seq counterparts. To ensure mathematically valid integration without losing platform-specific variance, we employed a Probabilistic Manifold Anchoring strategy^19^, which constrains the L1000 projection within the dimension-specific bounds of the RNA-seq prior. To overcome the inherent scarcity of experimental perturbation data, AetherCell harnesses the extensive prior knowledge encapsulated in large-scale foundation models (Fig. 1b). We integrated MolFormer^20^ to embed small molecules, fine-tuning it via Low-Rank Adaptation (LoRA)^21^ to adapt its chemical structure representations to specific biological contexts. Similarly, gene perturbations were encoded by fusing protein sequence embeddings from ESM-C^22,23^ with global interaction topology from the STRING PPI network^24^ via a Graph Neural Network (GNN). Finally, a cross-attention mechanism predicts the mechanism-specific latent transition vector (Δz) induced by these perturbations, which is superposed onto the baseline cell-state vector to model the perturbed state trajectory. This platform-agnostic, context-aware architecture establishes AetherCell as a foundational generative model capable of navigating the complex hierarchy from molecular interactions to different biological states, providing a robust base for diverse downstream applications including virtual transcriptome experiments, high-fidelity drug response prediction and precision drug repurposing (Fig. 1c).

We validated that this framework effectively bridges platform barriers while retaining biological signal. Uniform Manifold Approximation and Projection (UMAP) visualizations demonstrated that while samples initially clustered by platform, our anchoring strategy successfully aligned matched cell lines despite significant technological discrepancies (Fig. 1d). This unification was quantitatively confirmed by substantial improvements in ANOSIM^25^ and Silhouette^26^ scores (Fig. 1e). Finally, to verify that this unified latent space preserves essential biological information despite high-level compression, we quantitatively assessed reconstruction fidelity on held-out test sets. The model demonstrated robust signal preservation, achieving a consistent median Pearson Correlation Coefficient (PCC) of 0.95 for both the diverse RNA-seq corpus and the L1000 platform (Fig. 1f). Notably, this performance was consistent across distinct perturbations, including compound treatments, gene knockout (KO), knockdown (KD), and overexpression (OE) (Fig. 1f), confirming that AetherCell accurately captures biological variance and lays a solid foundation for downstream tasks.

### AetherCell resolves mechanism-specific signatures from systematic noise

We comprehensively evaluate whether AetherCell captures mechanism-specific signatures induced by perturbations. We first conducted a systematic benchmark on the L1000 dataset to evaluate AetherCell’s ability to predict transcriptomic shifts in unseen biological contexts. In the Unseen Cell scenario, AetherCell achieved a median Pearson Correlation Coefficient (PCC) of 0.82 on Differentially Expressed Genes (DEGs) (Fig. 2a). Notably, in the generally more challenging Unseen Compound scenario, the model achieved a median DEG PCC of 0.83, demonstrating its exceptional capacity to generalize to novel chemical entities (Fig. 2b). In both scenarios, AetherCell significantly outperformed state-of-the-art (SOTA) tool TranSiGen and other tools^27–30^ (*P* < 0.0001, Wilcoxon test).

**Fig. 2.**
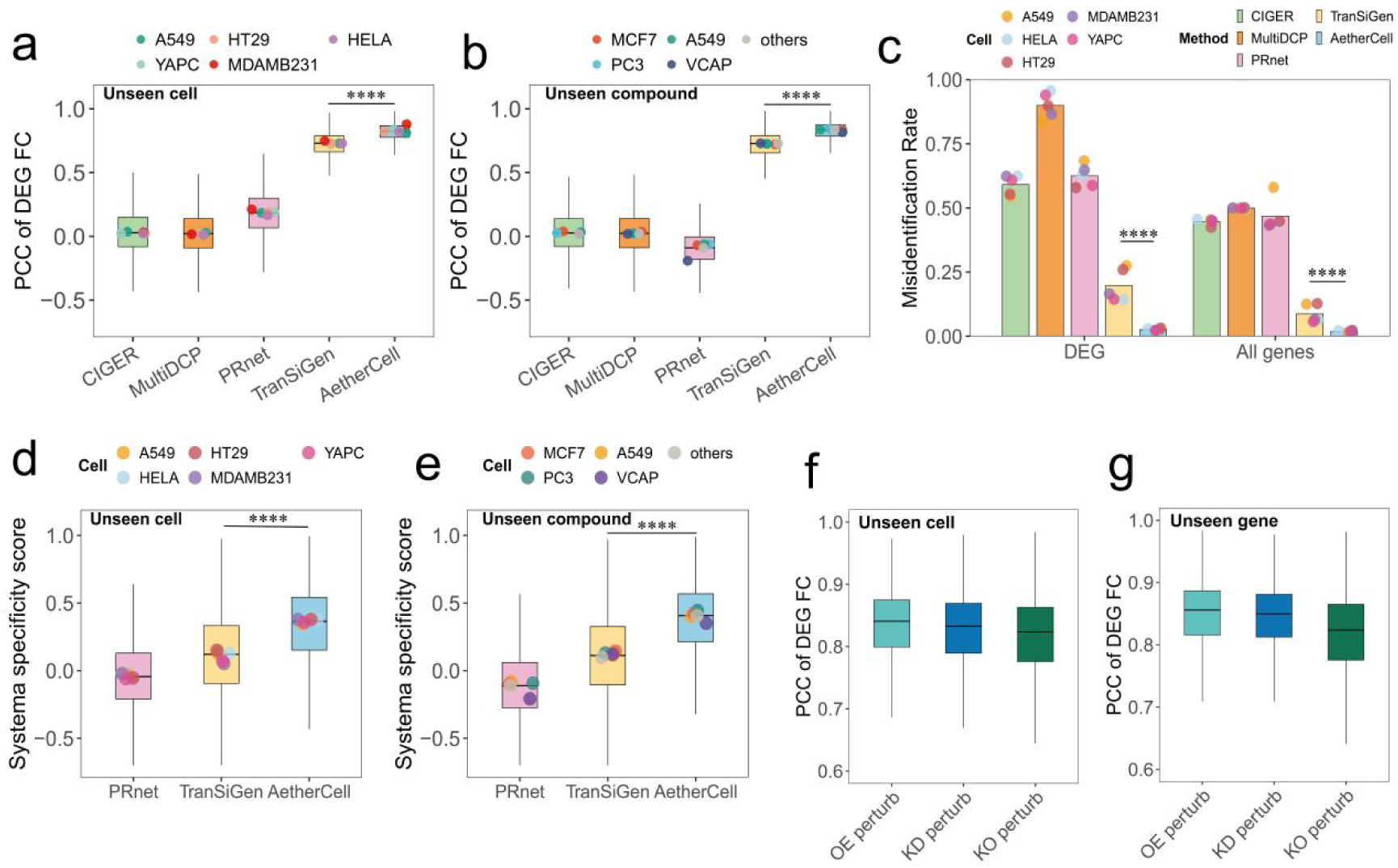
AetherCell demonstrates robust generalization and mechanism-specific prediction across chemical and genetic perturbations. a, b, Benchmarking perturbation prediction in unseen contexts. Box plots showing PCC of predicted vs. ground-truth log2FC (on DEGs, top 10% variance) for AetherCell versus baselines in strictly held-out **(a)** Unseen cell (*n* = 326,497) and **(b)** Unseen compound (*n* = 239,250) scenarios. **c,** Perturbation Misidentification Rate (*n* = 6,485); lower values indicate higher prediction tightness relative to random centroids. **d, e,** Systema Specificity Scores comparing AetherCell and baselines in **(d)** Unseen cell (*n* = 326,497) and **(e)** Unseen compound (*n* = 239,250) scenarios. **f, g,** Generalization to genetic perturbations (OE, KD, KO). Performance (PCC of DEG log2FC) evaluated in (f) Unseen Cell (OE: *n* = 17,081; KD: *n* = 55,136; KO: *n* = 58,756) and **(g)** Unseen Gene (OE: *n* = 11,979; KD: *n* = 49,036; KO: *n* = 36,264) settings. Box plots indicate median, quartiles, and 1.5× IQR. Statistical significance: two-sided Wilcoxon tests (**** *P* < 0.0001).

Relying solely on global correlations can be misleading. A central challenge in perturbation modeling is the dominance of “systematic noise”—generic, high-frequency transcriptomic signatures like cellular stress or metabolic shifts that appear across diverse treatments^15^. Because these features are statistically prevalent, standard models often succumb to “Mean-State Convergence”—a phenomenon we formally characterize as Type II Failure (Methods 2.2), where models regress to a statistical centroid that yields deceptively high global correlations but fails to distinguish the specific changes of the query perturbation. To strictly evaluate the resolution of mechanism-specific signals amidst this noise, we assessed the Perturbation Misidentification Rate^15^, a rigorous retrieval metric that quantifies the risk of a predicted profile being confused with other distinct perturbations. AetherCell attained a near-optimal (lowest) score of 0.03, significantly outperforming TranSiGen (0.20) and other representative tools which showed a high tendency toward profile misidentification (ranging from 0.59 to 0.90; Fig. 2c; *P* < 0.0001, Wilcoxon test).

We further utilized the Systema Specificity Score^15^, a robust metric designed to isolate “low-frequency” mechanistic drivers from the systematic stress baseline. For both Unseen Cell and Unseen Compound test sets, AetherCell achieved significantly higher scores of 0.37 (Fig. 2d) and 0.41 (Fig. 2e), respectively, significantly higher than other tools (TranSiGen: ∼ 0.12; PRnet: ∼ -0.04; P < 0.0001, Wilcoxon test). These metrics confirm that AetherCell’s predictions are not generic averages but remain tightly anchored to their specific ground-truth identities.

AetherCell’s ability to resolve these subtle signals is driven by a Specificity-Driven Generative framework designed to counteract identity dominance. Ablation studies confirmed that standard reconstruction objectives fail to disentangle specific MoA signals from background noise, resulting in a collapse toward generic stress centroids (Supplementary Fig. 3). Consequently, our multi-scale objective framework explicitly penalizes convergence to these generic backgrounds while strictly supervising the directional trajectory of high-variance ‘driver genes.’ This ensures that the generated transitions reflect the unique fingerprint of each perturbation rather than statistical averages.

This robustness extends to genetic perturbations, where the cross-attention module was specifically trained on diverse genetic intervention datasets. Under strictly held-out settings where target genes were not encountered during training, the model demonstrated balanced and high-fidelity performance across gene OE, KO, and KD tasks, with median DEG PCCs consistently exceeding 0.80, a comparable performance with compound perturbation (Fig. 2f-g). The ability of the model to resolve these specific transcriptomic signatures within a unified manifold suggests a robust capacity for cross-platform and cross-context generalization.

### Generalization to whole transcriptome and to complex tissue architectures

We reasoned that the platform-invariance of the AetherCell foundation model would enable a critical translational capability: the direct transfer of drug-induced regulatory logic from sparse, landmark-based screens to whole-transcriptomic, primary-mimetic environments without additional training. This addresses the fundamental barriers that have historically limited the utility of cell-line-based datasets.

We first evaluated AetherCell’s cross-platform generalization on independent RNA-seq datasets. Even though training perturbation data from the L1000 platform only measures a reduced representation of 978 landmark genes, AetherCell demonstrated robust performance on 744 paired RNA-seq datasets with compound perturbations. This super-resolution capability is underpinned by the Deep Backbone Decoder, which leverages the unified latent manifold to infer genome-wide expression changes from the compressed perturbation embeddings. The model achieved high predictive fidelity for both DEG fold changes and overall gene expression levels, significantly outperforming existing methodologies (Fig. 3a; *P* < 0.0001, Wilcoxon test). Notably, while TranSiGen exhibited relatively good performance on L1000 benchmarks, it demonstrated negligible predictive power for DEG identification when transferred to independent RNA-seq test sets.

**Fig. 3.**
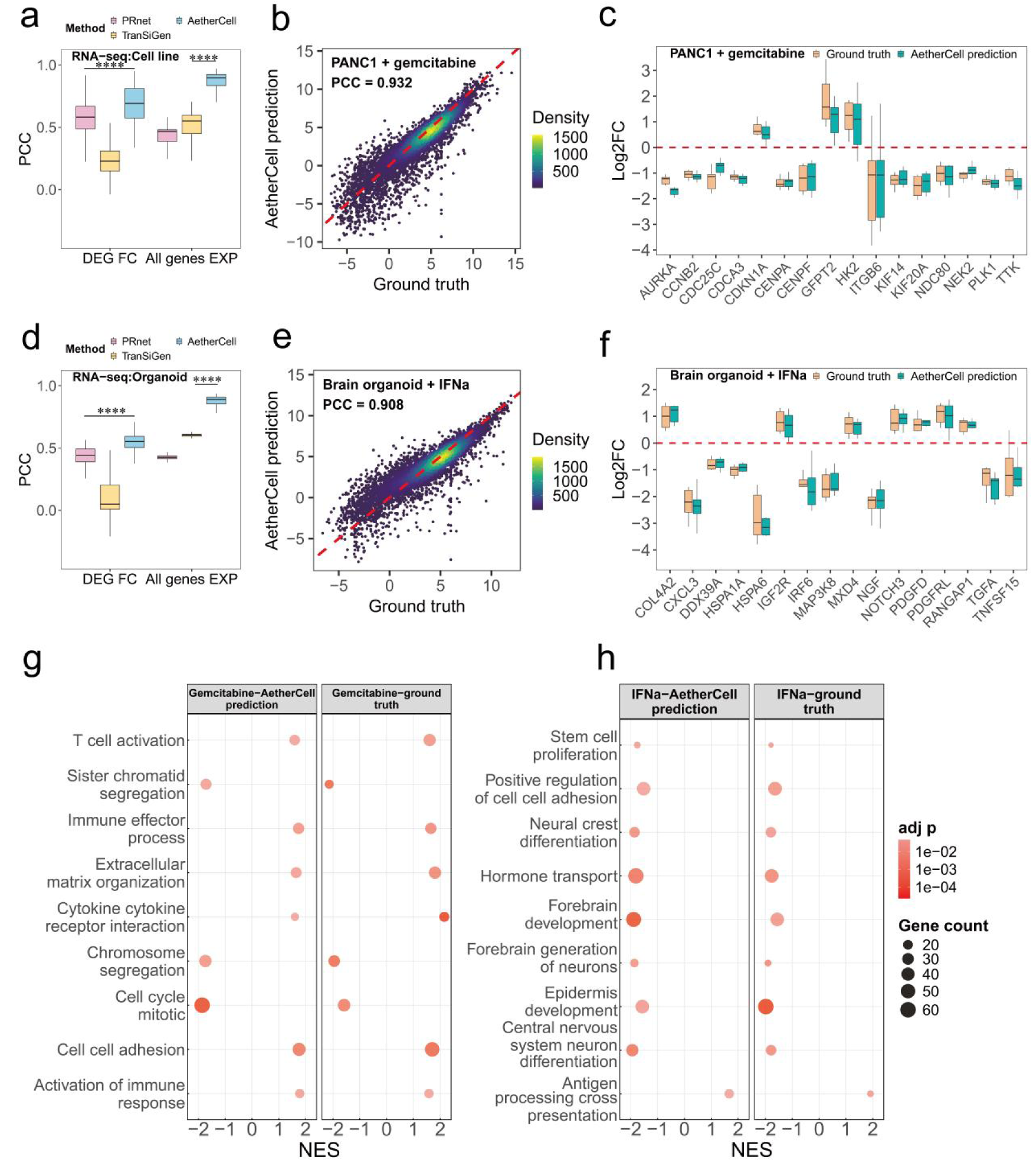
Cross-platform transferability and zero-shot generalization to complex biological systems. a, d, Benchmarking on independent (a) cell line (*n* = 658) and (d) organoid (*n* = 86) RNA-seq datasets. Box plots evaluate AetherCell versus baselines using PCC of DEG log2FC and absolute expression (All genes EXP). **b, e,** Global transcriptional reconstruction. Density scatter plots showing predicted vs. ground-truth expression for (b) Gemcitabine-treated PANC-1 cells and (e) IFN-treated brain organoids. **c, f,** Gene-wise fidelity. Box plots comparing ground-truth (beige) and predicted (teal) log2FC for top variant genes in (c) PANC-1 (*n* = 16) and (f) brain organoid (*n* = 12) scenarios. Red dashed lines indicate zero change. **g, h,** GSEA pathway comparison. Dot plots of Normalized Enrichment Scores (NES) for (g) Gemcitabine-treated PANC-1 and (h) IFN-treated organoids. Box plots indicate median, quartiles, and 1.5× IQR. Statistical significance: two-sided Wilcoxon tests (**** *P* < 0.0001).

To evaluate whether AetherCell can recover biologically interpretable mechanisms in a whole-transcriptomic environment, we examined the cellular response to gemcitabine^31^, a nucleoside analog that inhibits DNA synthesis, in the PANC-1 cell line as a representative validation. AetherCell demonstrated a high reconstruction fidelity of the overall gene expression profile (PCC = 0.932, Fig. 3b). Moreover, the model precisely captured the top differentially expressed genes, including critical cell cycle regulators such as AURKA, CCNB2, and CDC25C (Fig. 3c), showing high point-to-point correspondence with the experimental ground truth. Subsequent Gene Set Enrichment Analysis (GSEA) accurately recovered the drug’s underlying molecular mechanisms (Fig. 3g). Specifically, the predicted transcriptome mirrored the ground truth by correctly identifying the significant suppression of proliferation-associated pathways, such as DNA replication, DNA repair, and mitotic cell cycle processes, consistent with gemcitabine’s role as a DNA synthesis inhibitor. Simultaneously, the model successfully captured the upregulation of stress-response signaling, such as the activation of immune responses and response to tumor necrosis factor. These results confirm that the specificity-driven training objectives successfully disentangled transferable biological mechanisms from platform-specific noise.

To evaluate AetherCell’s robustness in primary-mimetic environments, we examined its performance on complex 3D organoid models, which represent a significant increase in biological heterogeneity over immortalized cell lines. By utilizing the organoid’s baseline transcriptomic profile as the context query in our cross-attention module, the model was able to adapt its perturbation predictions to these previously unseen biological contexts. We benchmarked this capability on 86 paired “Pre- vs. Post-perturbation” transcriptomic datasets spanning 12 distinct organoid models treated with 23 diverse chemical perturbations. Remarkably, despite being entirely naive to perturbation-induced signatures in organoids during training, AetherCell achieved high-fidelity reconstruction of the organoid-specific transcriptomic landscape. This zero-shot performance was comparable to results achieved on cell-line datasets (Fig. 3a) and significantly outperformed all existing methodologies (Fig. 3d; *P* < 0.0001, Wilcoxon test).

Beyond general benchmarks, we investigated whether AetherCell could capture the complex dynamics of viral pathogenesis and therapeutic intervention in HSV-infected brain organoids treated with IFNα. Despite never encountering the specific regulatory landscape of human neurodevelopment or viral infection during training, AetherCell reconstructed the perturbed state with high fidelity (median PCC = 0.908; Fig. 3e). In addition, it accurately predicted the upregulation and downregulation of important biomarkers^32^ (Fig. 3f). GSEA confirmed that the model captured the distinct biological shift induced by the treatment^33^ (Fig. 3h). Specifically, the predictions accurately recovered the activation of antiviral defense mechanisms while simultaneously capturing the suppression of neurodevelopmental and proliferative modules, effectively mirroring the tissue’s transition from a growth state to an immune-defense state. This translational consistency demonstrates that the unified latent space preserves the regulatory grammar required to generalize across disparate biological contexts.

### High-fidelity drug response prediction, patient stratification and clinical translation

Developing an accurate predictive model for drug response remains a primary challenge for both clinical diagnosis and drug development. We utilized perturbation-specific latent embeddings (Δz) derived from AetherCell to fine-tune a drug response model (AetherCell-RP) on drug sensitivity data from GDSC2^34^, a pharmacogenomic dataset providing standardized drug sensitivity profiles across hundreds of human cancer cell lines. Utilizing a 4:1 random split of the GDSC2 dataset for validation, AC-RP achieved a PCC of 0.92 (Fig. 4a) across 24,133 independent records (all *P* < 0.0001, Wilcoxon test). Even for unseen cell lines, our model achieved a PCC of 0.848 in predicting their drug response (Supplementary Fig. 10). Beyond regression accuracy, the model demonstrated high performance in binary stratification, distinguishing responders from non-responders with an AUROC of 0.944 and 0.982 on two independent large-scale datasets, GDSC^34^ (n=37,662) and PRISM^35^ (n=52,717) (Fig. 4b). This substantial performance confirms that explicitly modeling the latent transition provides a high-resolution signal that captures complex, non-linear interactions.

**Fig. 4.**
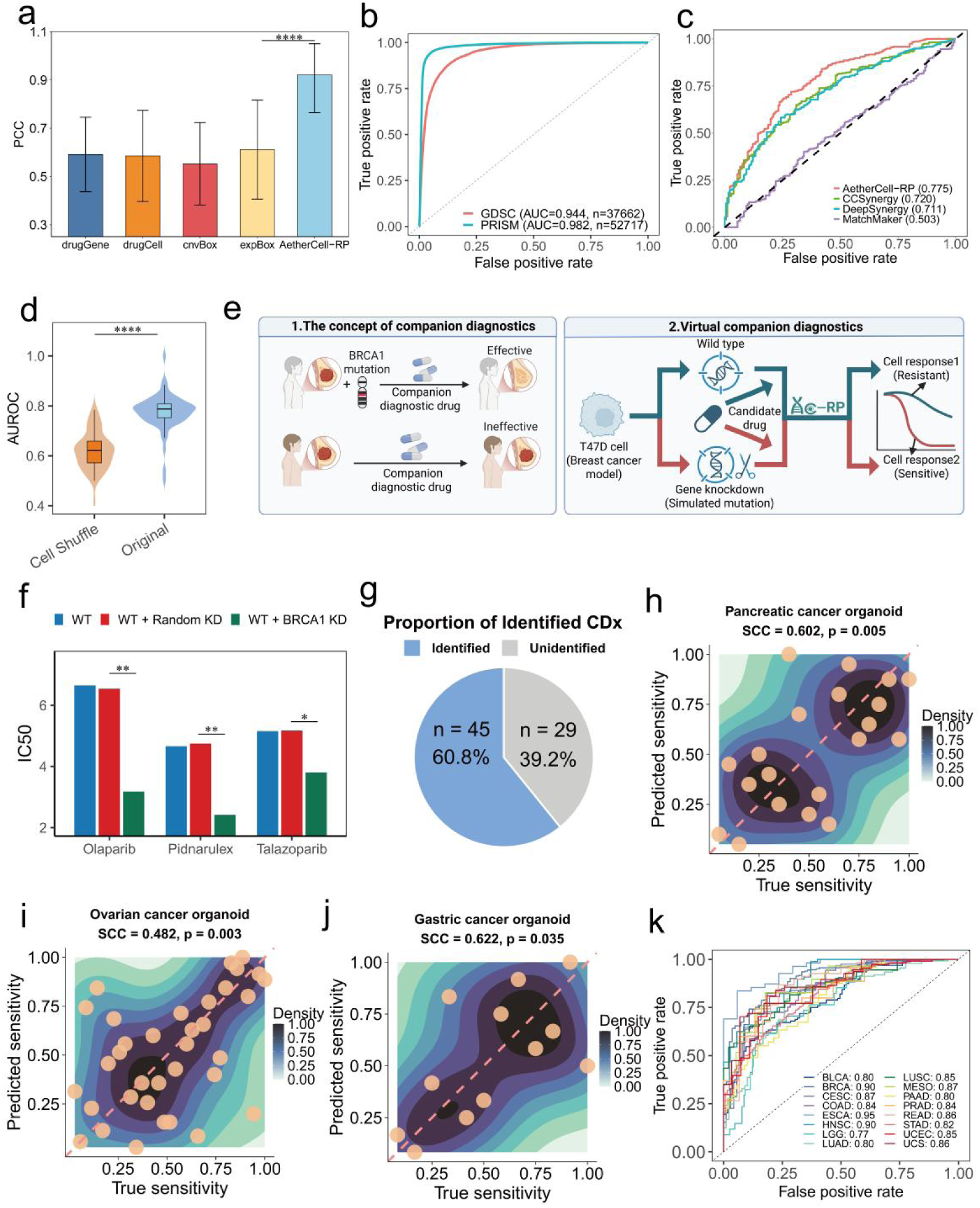
Mechanism-driven drug response prediction and translational generalization to clinical phenotypes. a, Dynamic perturbation-based sensitivity prediction. Bar plot of PCC for predicted vs. actual IC50 (*n* = 24,133) comparing AetherCell-RP to genomic baselines. Data are presented as mean ± *s*.*d*. Significance: two-sided Wilcoxon (**** *P* < 0.0001 vs. expBox). **b**, Classification of drug sensitivity. ROC curves for GDSC (*n* = 37,662) and PRISM (*n* = 52,717) datasets. **c, d,** Synergy prediction. (c) ROC curves versus baseline models. (d) Violin plots comparing AUROC of AetherCell-RP vs. Cell Shuffle control (two-sided Wilcoxon, **** *P* < 0.0001). **e, f, g,** Virtual Companion Diagnostics (CDx). (e) Schematic workflow. (f) Synthetic lethality assessment for PARP and rRNA inhibitors. Bar plots show predicted IC50 for Wild Type (WT), Random KD, and BRCA1 KD. Significance: random permutation test (*n* = 1,503; \**P* < 0.05, \*\**P* < 0.01). (g) Pie chart of identified clinical biomarkers (*n* = 45 identified, *n* = 29 unidentified). **h, i, j,** Zero-shot generalization to PDOs. Density scatter plots of predicted vs. ground-truth IC50 (SCC) in (h) pancreatic (*n* = 20), (i) ovarian (*n* = 35), and (j) gastric (*n* = 12) organoids. **k,** Clinical response prediction. ROC curves for TCGA patient cohorts (*n* = 2,966). Detailed descriptions of the cancer types corresponding to the TCGA codes are provided in Supplementary Table. 2. Box plots indicate median, quartiles, and 1.5× IQR. Error bars represent mean.

We further extended this capability to predict combinatorial drug effects by fine-tuning AC-RP on drug synergy data from DrugComb^36^. Our model achieved SOTA performance in predicting drug synergy with an AUROC of 0.775 (Fig. 4c), outperforming three drug synergy prediction tools^37–39^. A “Cell Shuffle” control experiment, where cell-line identity labels were randomized, showed a significant performance reduction compared to the unshuffled (*P* < 0.0001, Wilcoxon test), confirming that the model leverages precise, cell-type-specific transcriptomic dependencies rather than memorizing global drug potencies (Fig. 4d).

Beyond predictive accuracy, AetherCell enables mechanism-driven *in silico* screening for Companion Diagnostics (CDx)—clinical biomarkers that identify patient subpopulations most likely to benefit from specific therapeutic interventions. Functioning as a “virtual laboratory,” the model simulates genetic variations to assess their impact on drug sensitivity via a counterfactual inference framework (Fig. 4e; Methods). By using gene knockdowns to proxy loss-of-function mutations, AetherCell-RP predicts differential drug sensitivity (IC50) for wild-type versus gene-deficient states. As a proof-of-concept for synthetic lethality and genotype-specific sensitivity, we simulated the effect of BRCA1 knockdown on responses to three distinct therapeutic agents: the PARP inhibitors olaparib and talazoparib^40^, and the rRNA synthesis inhibitor pidnarulex^41^. Despite these drugs belonging to different therapeutic classes with different biological mechanisms, AetherCell-RP correctly predicted that BRCA1-deficient states drastically sensitize cells to all three drugs compared to wild-type or random-knockdown controls (*P* = 0.005, 0.003 and 0.018, respectively, random permutation testing; Fig. 4f). Expanding this analysis to 74 clinically validated drug-gene pairs from CIViC (Clinical Interpretation of Variants in Cancer)^42^, the model successfully recovered 60.81% of known associations (Fig. 4g). This robust alignment with established clinical evidence underscores the model’s potential as a reliable virtual screening engine, capable of systematically prioritizing actionable biomarkers to guide precision oncology strategies.

In recent years, Patient-Derived Organoids (PDOs) have emerged as a preeminent platform for functional precision oncology—faithfully recapitulating the 3D architecture, genomic landscape, and histological complexity of primary tumors. However, their widespread clinical adoption remains constrained by high operational costs, lengthy establishment timeframes, and limited scalability compared to traditional *in vitro* assays. To bridge the gap, we leveraged AetherCell’s inherent platform-invariance to evaluate zero-shot generalization across complex, biologically heterogeneous models. Remarkably, without any exposure to primary tissue data during training, AetherCell’s predictions in PDOs demonstrated significant correlations with experimental ground truth in pancreatic^43^, ovarian^44^, and gastric cancers^45^ (*P* = 0.005, 0.003 and 0.035, respectively, Spearman correlation test; Fig. 4h-j). This high-fidelity, zero-shot generalization establishes AetherCell as a ‘virtual laboratory’ capable of scaling the biological insights of 3D architecture to large-scale patient cohorts where physical organoid establishment is unfeasible.

This translational fidelity further extended to the direct prediction of real-world clinical drug responses in large-scale patient cohorts from The Cancer Genome Atlas (TCGA). The dataset comprised ∼3,000 samples across 17 cancer types (minimum n = 50), with testicular cancer excluded due to extreme class imbalance between responders and non-responders (Supplementary Fig. 13). Despite inherent clinical heterogeneity, AetherCell’s foundational manifold required only minimal calibration via a histology-aware adaptive transfer framework (Methods 9). By freezing the pre-trained encoder and fine-tuning lightweight adapters (10,000 parameters) to integrate clinical covariates, the model accurately stratified responders from non-responders in “Unseen Patient” settings, with AUROCs exceeding 0.80 for 15 out of 16 cancer types (Fig. 4k). This validates the transferability of our foundation model to complex clinical environments, demonstrating that it can effectively guide personalized treatment decisions even when patient data is scarce.

Collectively, these results establish that AetherCell transcends the limitations of its cell-line-based training data, utilizing its unified latent space as a robust foundational manifold to achieve successful translation from high-throughput screening to diverse human physiological contexts.

### Uncovering novel therapeutic interventions via adaptive phenotype-knowledge integration

Current drug repurposing strategies face a fundamental “granularity-generality paradox”: knowledge graph (KG)-based methods excel at macro-scale indication prediction but lack the resolution to capture cell-type specific heterogeneity, whereas transcriptomics-based methods capture micro-scale phenotypic reversals but struggle to generalize across systemic diseases due to signal sparsity. To this end, we developed AetherCell-DR (drug repurposing), a Phenotype-Knowledge Mixture of Experts (PK-MoE) system designed to adaptively route inference based on the specific biological context (Fig. 5a). Following the principle of orthogonal complementarity, the model decomposes drug efficacy into two distinct pathways: physical phenotypic reversal, handled by a Transcriptomic Expert operating on the platform-invariant Δ*z* embeddings of AetherCell, and target/pathway blockade, handled by a Knowledge Expert based on biomedical knowledge graph (Methods 10). A context-aware gating network acts as a dynamic “triage system,” assigning sample-specific weights (*w_trans_, w_know_*) to each expert by evaluating the strength of the predicted transcriptomic reversal signal relative to the local connectivity density of the drug-disease pair within the knowledge graph.

**Fig. 5.**
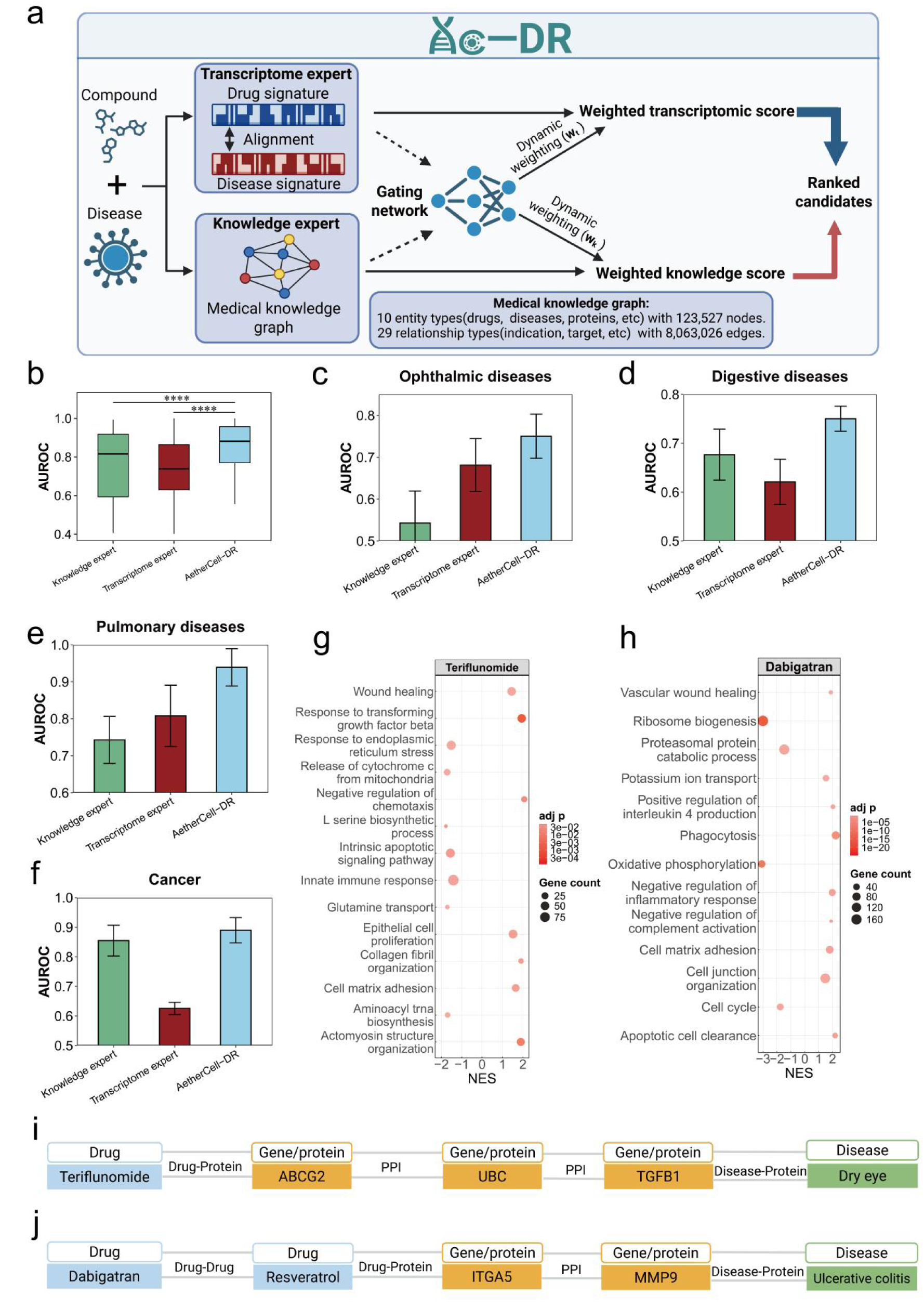
Adaptive drug repurposing via Phenotype-Knowledge Mixture of Experts (PK-MoE). a, Architecture of the AetherCell-DR framework. A dual-expert system fusing macro-scale associations (Knowledge Expert) and micro-scale expression changes (Transcriptomic Expert) via a Context-Aware Gating Network. **b,** Performance evaluation on macro-scale clinical indications. Box plots of AUROC scores (*n* = 196) comparing AetherCell-DR to individual experts. Significance: two-sided paired Wilcoxon signed-rank tests (**** *P* < 0.0001). **c–f,** Comparisons across diverse disease cohorts (*n* = 20). Bar plots showing performance for (c) Ophthalmic, (d) Digestive, (e) Pulmonary, and (f) Cancer diseases. Specific diseases included in each category are listed in Supplementary Table 3. Error bars represent mean. **g–j,** Multi-modal interpretability. (g, h) Transcriptomic GSEA showing phenotypic reversal for (g) teriflunomide and (h) dabigatran. (i, j) Knowledge reasoning paths connecting (i) teriflunomide to DED via ABCG2-TGFB1 and (j) dabigatran to UC via MMP9. Box plots indicate median, quartiles, and 1.5× IQR.

We tested our model in predicting FDA-approved drugs for 196 systemic diseases, the same task evaluated in the study of the SOTA model, TxGNN^46^. The integrated MoE system achieved a median AUROC of 0.88 (Fig. 5b), significantly outperforming either standalone expert (*P* < 0.0001, Wilcoxon test), where the Transcriptomic Expert obtained an AUROC of 0.72 and Knowledge Expert (TxGNN) obtained an AUROC of 0.81. Crucially, we validated that the MoE resolves the granularity-generality paradox through autonomous “division of labor”: while the Knowledge Expert collapsed to near-random performance (median AUROC = 0.52) on fine-grained cell line sensitivity tasks, both the Transcriptomic Expert and the integrated MoE maintained robust predictive accuracy (Supplementary Figure. 14). Diverse therapeutic areas confirmed the robustness of AC-DR, including the ophthalmic (AUC = 0.75), digestive (AUC = 0.75), pulmonary (AUC = 0.94), and cancer (AUC = 0.89) cohorts (Fig. 5c-f). These results confirm that the MoE architecture successfully harmonizes systemic priors with transcriptomic specificity, enabling the model to generalize effectively across distinct organ systems and capture complex disease-drug associations.

To demonstrate the translational utility of this framework for uncovering novel therapeutics, we applied AetherCell-DR to Dry eye disease (DED), a highly prevalent multifactorial ocular condition that imposes a substantial socioeconomic burden and significantly impairs the quality of life and visual function of around 25% of the adult population worldwide^47^. To prevent information leakage, all eye-disease-related terms were removed from the knowledge graph during the training phase. Among the top 10 candidates identified by AC-DR, two FDA-approved or standard-of-care treatments (including the recently approved varenicline), three drugs currently under clinical investigation or with established off-label clinical evidence, and three agents supported by preclinical studies (Supplementary Table. 4). This represents a substantial hit rate in the discovery of translatable therapeutic options. Surprisingly, the model prioritized teriflunomide^48^, a DHODH inhibitor conventionally used for multiple sclerosis, even though neither teriflunomide nor its target has been previously associated with DED in the literature. Attention-weighted path analysis provided a convergent rationale: The Transcriptomic Expert predicted that teriflunomide significantly upregulates “wound healing” and “cell matrix adhesion” pathways, suggesting a role in corneal epithelial repair (Fig. 5g). Simultaneously, the Knowledge Expert identified a mechanistic path linking teriflunomide to the ABCG2 transporter and the UBC ubiquitin gene, which subsequently regulates the TGF1 axis—a critical modulator of ocular inflammation (Fig. 5i). This multi-modal convergence suggests that teriflunomide may function as a dual-action therapeutic, promoting structural restoration while dampening cytokine-mediated inflammation.

We also utilized the model to screen for novel candidates in Ulcerative Colitis (UC), a chronic, relapsing inflammatory bowel disease of the colon and rectum that has become a growing global health burden^49^. Evaluation of the top 10 repurposed candidates reveals exceptional clinical alignment: our model successfully recovered eight FDA-approved or standard-of-care agents, and one drug currently supported by clinical trial evidence for IBD, demonstrating the model’s reliability in capturing established therapeutic efficacy (Supplementary Table. 5). Notably, the thrombin inhibitor dabigatran^50^ was listed in the top-hit list—an unexpected repositioning from anticoagulation to gastrointestinal medicine. Transcriptomic analysis predicted that dabigatran would reverse the UC-associated inflammatory signature by upregulating “Vascular wound healing” and “Cell junction organization,” potentially accelerating the repair of the intestinal mucosal barrier (Fig. 5h). Complementing this, the Knowledge Expert constructed a mechanistic reasoning chain linking dabigatran to the modulation of MMP9 (Matrix Metallopeptidase 9) via ITGA5/Resveratrol-like interaction nodes (Fig. 5j). Given that MMP9 is a primary driver of tissue degradation and barrier breakdown in UC, this suggests a plausible mechanism where dabigatran exerts protective effects by stabilizing the tissue remodeling microenvironment. These case studies demonstrate that AC-DR goes beyond simple association prediction, serving as an AI framework for the hypothesis-driven generation of novel treatments.

### *In vivo* validation of novel therapeutic interventions

Given the unexpected nature of the computational predictions, we performed *in vivo* validation of teriflunomide for DED and dabigatran for UC. We first employed a well-established benzalkonium chloride (BAC)-induced mouse model^51^ to evaluate the therapeutic potential of teriflunomide (Fig. 6a). We included loteprednol etabonate^52^, an FDA-approved DED treatment, as a positive control. Clinical assessment via slit-lamp microscopy showed teriflunomide treatment reduced corneal opacity compared to the vehicle group and matched the performance of the clinical positive control (Fig. 6b). Investigation of the conjunctival microenvironment via Periodic Acid-Schiff (PAS) staining confirmed a potent restorative effect, with teriflunomide treatment significantly increasing goblet cell density (*P* = 0.0005, ANOVA test; Fig. 6c and d, Supplementary Fig. 17). Furthermore, fluorescein staining demonstrated the effective healing of corneal epithelial defects (Fig. 6e). At the histological level, teriflunomide successfully reconstructed the corneal architecture, restoring epithelial thickness from a damaged state (∼20 *μm*) to a healthy, stratified morphology (∼38 *μm*), achieving parity with the positive control (Fig. 6f; representative images in Supplementary Fig. 16). This structural restoration was functionally corroborated by the tear secretion test, which showed a significant reversal of dry eye symptoms that was indistinguishable from the efficacy of loteprednol etabonate (*P* < 0.0001, ANOVA test, Fig. 6g). Collectively, these findings confirm that teriflunomide acts as a potent and effective protective agent for DED, providing therapeutic outcomes equivalent to current standard-of-care treatments.

**Fig. 6.**
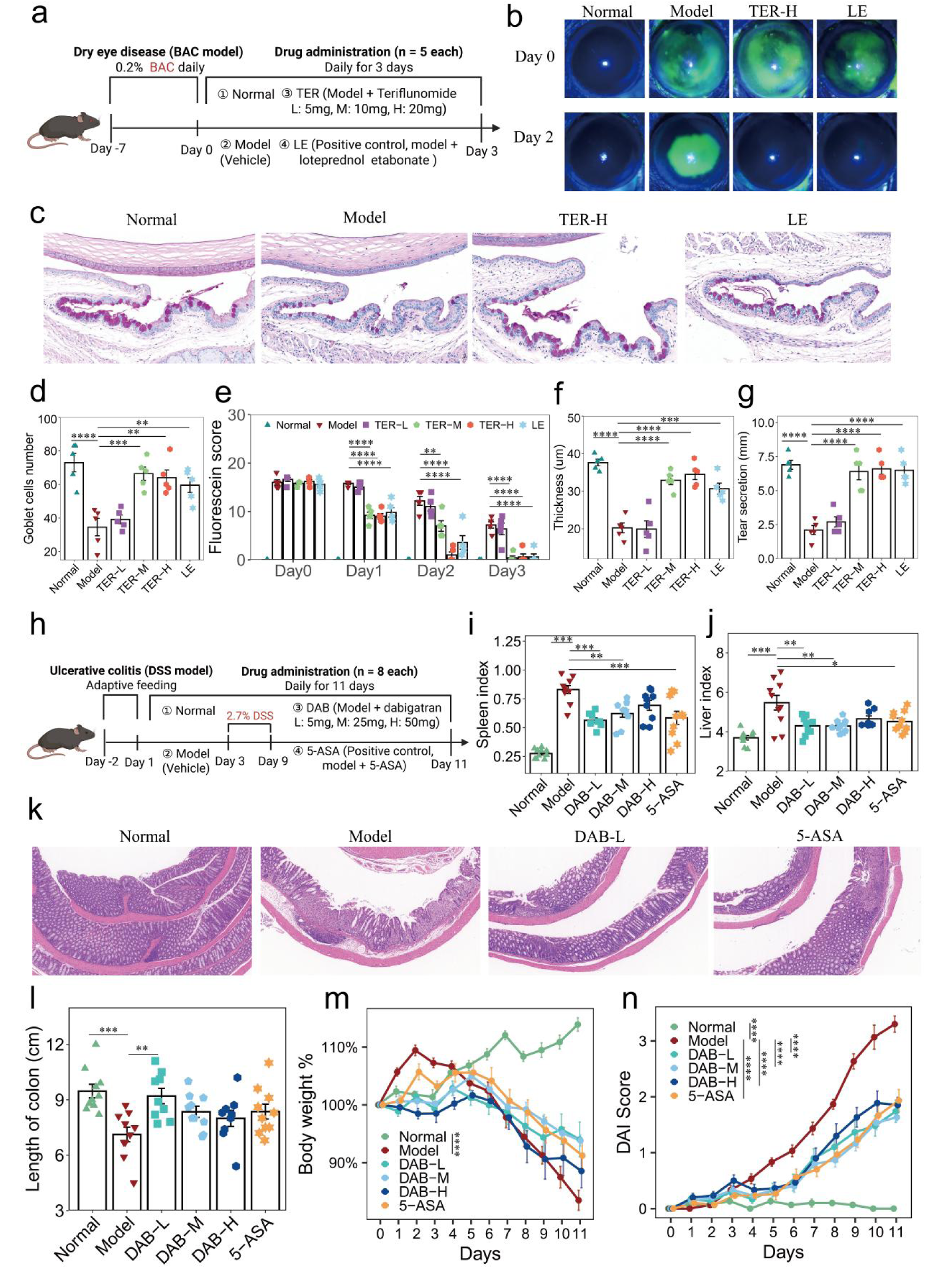
In vivo validation of AI-prioritized first-in-class therapeutic candidates across distinct pathologies. a-g, Validation in Dry Eye Disease (DED) model (*n* = 5). (a) Experimental design using BAC-induced model treated with teriflunomide vs. loteprednol etabonate. (b) Slit-lamp and (c) PAS staining images. Quantitative assessment of (d) goblet cell density, (e) fluorescein scores, (f) epithelial thickness (*μm*), and (g) tear secretion (mm). **h–n,** Validation in Ulcerative Colitis (UC) model (*n* = 8). (h) Experimental design using DSS-induced model treated with dabigatran vs. 5-ASA. (i, j) Systemic inflammation via (i) spleen and (j) liver indices. (k) H&E staining. (l) Colon length. (m, n) Longitudinal monitoring of (m) body weight and (n) Disease Activity Index (DAI). Error bars represent mean. Significance: one-way or two-way ANOVA with Tukey’s test (∗ ∗ ∗ ∗ *P* < 0.0001; ∗ ∗ ∗ 0.0001 < *P* ≤ 0.001; ∗ ∗ 0.001 < *P* ≤ 0.01; ∗ 0.01 < *P* ≤ 0.05).

We next assessed dabigatran, a direct thrombin inhibitor identified by AC-DR as a novel candidate for UC. We employed 5-Aminosalicylic Acid (5-ASA)^53^, an FDA-approved clinical standard for UC, as the positive control (Fig. 6h). Treatment with dabigatran effectively mitigated systemic inflammation, significantly reversing DSS-induced^54^ hepatosplenomegaly as indicated by normalized spleen and liver indices, with efficacy comparable to that of 5-ASA (Fig. 6i–j). At the tissue level, histopathological analysis confirmed that dabigatran treatment preserved the mucosal barrier structure and minimized inflammatory infiltration compared to the vehicle group (Fig. 6k, Supplementary Fig. 19). Macroscopically, this structural protection translated to the maintenance of colonic integrity; dabigatran exhibited significant protection against colon shortening (*P* = 0.005, ANOVA test), matching the performance of the positive control (Fig. 6l, Supplementary Fig. 18). Furthermore, dabigatran treatment alleviated the systemic impact of colitis, attenuating body weight loss throughout the acute phase to a degree indistinguishable from 5-ASA (Fig. 6m). Longitudinal monitoring of the Disease Activity Index (DAI) consistently confirmed these benefits, demonstrating that dabigatran delayed symptom onset and significantly reduced overall disease severity, achieving parity with the established clinical standard (Fig. 6n).Notably, dabigatran treatment did not exacerbate intestinal hemorrhage but instead led to a reduction in fecal occult blood compared to the untreated model group (Supplementary Fig. 20).

Collectively, these *in vivo* studies confirm that AC-DR successfully identifies high-value therapeutic candidates. The success across two physiologically distinct disease models underscores the framework’s robustness. These results establish AetherCell as a powerful engine for uncovering potential high-value therapeutic opportunities.

## Discussion

The development of AetherCell represents a fundamental shift in virtual cell modeling, transitioning from task-specific architectures to a foundational, platform-aligned framework that addresses the core “data-utility paradox” of modern transcriptomics. By bridging the dense perturbational coverage of landmark assays with the rich contextual diversity of clinical RNA-seq, we provide a “universal coordinate system” that enables the direct translation of molecular insights into human physiological environments. This capability is particularly timely given the global regulatory pivot toward Non-Animal Methods (NAMs), such as the U.S. FDA Modernization Act 3.0.

An important contribution of this work is the formal characterization and solution of “Mean-State Convergence” (Type II Failure). In generative biology, models frequently achieve deceptively high global correlations by reproducing high-frequency “generic” responses—such as cellular stress, apoptosis, and metabolic shifts—that are statistically prevalent across diverse perturbations. Such correlations can be misleading; a model can appear statistically successful while failing to resolve the unique mechanistic driver of a specific intervention. By implementing a specificity-driven learning framework, AetherCell explicitly suppresses these non-specific stress centroids. The significantly lower Perturbation Misidentification Rate and higher Systema Specificity Scores achieved by AetherCell suggest that “accuracy” in perturbation modeling must be redefined to prioritize mechanistic resolution over generic statistical similarity, ensuring that virtual experiments recover true biological transitions rather than statistical averages.

The robustness of this foundational manifold is evidenced by its generalization across disparate biological scales, from 3D brain organoids to clinical cohorts. The model’s ability to accurately predict the suppression of neurodevelopmental modules in HSV-infected organoids—without having encountered viral dynamics during training—suggests an internalized “universal grammar” of gene regulation. This transcriptomic fidelity translates directly into functional utility through AC-RP, which achieves high-resolution prediction of drug sensitivity and genotype-linked vulnerabilities (Companion Diagnostics), in unseen cell lines, unseen patient-derived organoids (PDOs) and real-world TCGA cohorts. The achievement of AUROCs exceeding 0.80 across diverse cancer types confirms that the latent transitions (Δz) learned by AetherCell capture clinically actionable features rather than platform-specific artifacts, providing a scalable alternative to physical 3D culture establishment.

To resolve the “granularity-generality paradox” in therapeutic discovery, we introduced AC-DR, a Phenotype-Knowledge Mixture of Experts (PK-MoE) system. By adaptively routing inference between a Transcriptomic Expert (focused on molecular reversal) and a Knowledge Expert (focused on systemic priors), the model overcomes the limitations of either modality in isolation. The successful *in vivo* validation of teriflunomide for Dry Eye Disease and dabigatran for Ulcerative Colitis—two non-obvious repositioning candidates—demonstrates that this hybrid approach can surface high-value therapeutic hypotheses that bypass traditional discovery silos.

Despite these advancements, several limitations warrant future investigation. First, while AetherCell effectively leverages the scale of bulk RNA-seq to bridge perturbations with clinical contexts, the current iteration does not explicitly model intra-tissue heterogeneity. Transitioning toward single-cell transcriptomic foundations would allow the model to resolve how interventions modulate specific sub-populations within a niche. Second, while our cross-attention module is context-aware and effectively integrates cellular baseline states, it currently treats perturbations as static snapshots; integrating temporal dynamics will be essential for capturing the time-dependent non-linear trajectories of drug response. Finally, while transcriptomics is a powerful proxy, it represents only one layer of the central dogma; a truly comprehensive “Virtual Cell” will require the integration of proteomics and metabolomics into a unified multi-omic foundation.

In conclusion, AetherCell moves the field closer to the realization of a “Virtual Laboratory.” By providing a platform-invariant manifold that links molecular interventions to human physiological states, we establish a framework for the systematic, human-centric prioritization of therapeutics. As biological data scales in both volume and complexity, foundation models like AetherCell will be indispensable for transforming fragmented datasets into a cohesive, predictive, and actionable understanding of human health and disease.

## Author contributions

Z.X. conceived, designed and supervised the project. WY.L. developed deep learning models, including AetherCell, AetherCell-RP, and AetherCell-DR. Y.C. collected and preprocessed the datasets, evaluated model performance, and analyzed the data. ZY.P. conducted *in vivo* DED experiments. L.X. conducted in *vivo* UC experiments. D.W. designed and supervised the UC experiments. WY.L., Z.X. and Y.C. wrote the manuscript. All authors read and approved the manuscript.

## Data and code availability

The L1000 gene expression profiles used for model pre-training are publicly available at the CMap LINCS resource (https://clue.io/). Publicly available RNA-seq datasets used for model testing were derived from the Gene Expression Omnibus (GEO, https://www.ncbi.nlm.nih.gov/geo/). Detailed accession numbers and sample metadata for disease-related datasets are provided in Supplementary Data 1, and those for drug perturbation datasets are listed in Supplementary Data 2, and the source metadata for GEO datasets used in pre-training are summarized in Supplementary Data 3. Datasets for downstream benchmarking and validation tasks were obtained from the following public repositories: Drug sensitivity data: Sourced from the Genomics of Drug Sensitivity in Cancer (GDSC, https://www.cancerrxgene.org/) and the PRISM Repurposing dataset (DepMap, https://depmap.org/portal/prism/). Drug synergy data: Sourced from the DrugComb portal (http://drugcombdb.denglab.org/main). Clinical drug response data: Sourced from The Cancer Genome Atlas (TCGA, accessible via the GDC Data Portal at https://portal.gdc.cancer.gov/). Companion diagnostic (CDx) data: Sourced from the Clinical Interpretations of Variants in Cancer (CiVIC) database (https://civicdb.org/). All other data supporting the findings of this study are available within the article and its Supplementary Information files. Source data are provided with this paper.Code availability The source code for AetherCell is available on GitHub at (https://github.com/Wenyuan-AI4science/AetherCell), and training dataset can be accessed via zenodo (https://zenodo.org/records/18295255).

